# A common mechanism and multiple advantages of pigment loss underlie the convergent evolution of albinism in cave animals

**DOI:** 10.1101/2025.07.14.664374

**Authors:** Helena Bilandžija, Kenneth J. Renner, Helena Ćetković, William R. Jeffery

## Abstract

The loss of pigmentation is a hallmark adaptation of cave-dwelling animals but the underlying mechanisms are poorly understood. This study investigates the cellular and biochemical basis of albinism, the loss of melanin pigment, which convergently evolved across a wide range of cave animals. Is albinism caused by the interruption of pigment synthesis or elimination of the pigment cell lineage? The results suggest that albinism evolved by a common mechanism, a block in the first step of the melanin biosynthesis pathway, the conversion of L-tyrosine to L-DOPA, in diverse albino cave animals ranging from annelids and mollusks to vertebrates. Pigment cells are conserved in all tested albino cave species and distributed in patterns resembling pigmentation in their close surface relatives. The cells capable of melanin synthesis when provided with L-DOPA substrate were detected at the sites of injuries in a cave annelid, cave mollusk, and cave teleost, suggesting roles in innate immunity during tissue repair, which may explain why pigment cells are conserved despite the loss of pigment production. Next, we focused on the longstanding issue of the evolutionary forces involved in the regression of pigmentation in cave animals and explored hypotheses of why losing pigmentation might be adaptive in cave environments. First, we tested the hypothesis that pigment regression may conserve energy, which is important for survival in the food-limited cave environment. Our results indicate that melanin synthesis has a marked energetic cost in the cavefish, *Astyanax mexicanus*. Lastly, we show that the disruption of melanin synthesis at its first biosynthetic step is correlated with increased dopamine levels in multiple depigmented cavefish populations, including populations that evolved albinism independently. This result supports a widespread tradeoff between the melanin and catecholamine synthesis pathways in *A. mexicanus*. We conclude that the interruption of the conversion of L-tyrosine to L-DOPA in the synthesis of melanin confers multiple advantages that could serve as targets of natural selection, supporting an adaptive hypothesis for the evolution of albinism in cave animals.

## Introduction

Convergent evolution is widespread, occurring not only between phyla or kingdoms but even across domains (Kaiser et al, 2011, Suzuki & Numata, 2014). Classic examples include the evolution of bioluminescence, carnivory and multicellularity across kingdoms, camera-type eyes in vertebrates and cephalopods, and antifreeze proteins in Arctic and Antarctic fishes. It is well established that convergence arises when divergent lineages adapt to similar environmental pressures and ecological niches. However, it remains far less clear whether convergent traits result from shared or distinct developmental, molecular, and evolutionary pathways. Uncovering the mechanisms underlying convergence may provide critical insights into the constraints and predictability of evolution (Mas et al., 2020). At a time of accelerating environmental change, the ability to predict evolutionary outcomes is essential for conservation and managing future biodiversity (Nosil et al., 2020).

The loss of pigmentation is one of the hallmarks of cave-adapted animals and has evolved in diverse organisms ranging from planaria to vertebrates, regardless of their type of pigment. As such, pigment loss in cave animals is one of the finest examples of convergent evolution in nature. In cave forms of the teleost *Astyanax mexicanus* and two species of cave planthoppers, melanin biosynthesis is blocked at the first step of the pathway (McCauley et al., 2004; Bilandžija et al., 2012). However, the mechanisms responsible for albinism in all other cave-adapted species remain unknown. Are the underlying causes for convergent loss of melanin the same or different in divergent cave animals?

Melanin is one of the most common dark pigments in nature and found in plants, fungi, and different animal phyla (Riley, 1997). Melanin biosynthesis begins with the amino acid L-tyrosine, which is obtained from the diet or synthesized by hydroxylation of L-phenylalanine. L-tyrosine is first converted into L-3,4-dihydroxyphenylalanine (L-DOPA), which is the rate-limiting step in the pathway. L-DOPA is then oxidized to dopaquinone, which is a critical branch point in the pathway. If dopaquinone undergoes further enzymatic reactions, this leads to the production of eumelanin. This pathway involves several enzymes and intermediate molecules, such as 5,6-dihydroxyindole-2-carboxylic acid (DHICA) and 5,6-dihydroxyindole (DHI). In vertebrates, melanin is synthesized in specialized lysosome-derived melanosomes, which are located in melanin-containing pigment cells called melanophores (reviewed in Slominski et al., 2012).

The specific enzymes, intermediates, and regulatory mechanisms involved in melanin synthesis can vary among different animal species. Insects convert L-tyrosine to L-DOPA using tyrosine hydroxylase, whereas vertebrates and many other organisms catalyze this reaction using tyrosinase (Sugumaran and Barek, 2016, Vavricka et al., 2014). Conversely, tyrosine hydroxylase catalyzes the synthesis of the catecholamine (CAT) dopamine by utilizing L-tyrosine substrate in vertebrates. CATs are important for various physiological and neurological processes, including serving as neurotransmitters (Fernstrom and Fernstrom, 2007).

The evolutionary drivers of regressed traits including pigmentation in cave animals, whether primarily driven by neutral processes and genetic drift or by natural selection, have been the subject of persistent and often contentious debate (reviewed in Culver et al., 2023). Obviously, the functions of pigmentation, including visual processes, protection from UV light, behavioral displays, mimicry, and camouflage (Sugumaran, 2002; Protas and Patel, 2008), are useless in the absolute darkness of caves. Charles Darwin, in his revolutionizing book on natural selection, suggested that regression in cave animals could be a consequence of disuse (Darwin, 1872). Indeed, one cannot imagine that negative selection against spontaneous mutations that result in albinism would exist in dark caves, whereas it is reasonable that albinos would be targets of predation or less desirable sexual partners on the surface. Thus, one hypothesis to explain the regression of pigmentation in albino cave animals has been neutral mutation and genetic drift in which purifying selection for mutations in the pigmentation pathways is presumed to be lacking (Juan et al., 2010; Wilkens and Strecker, 2017). However, it is difficult to reconcile the slow and stochastic progress of genetic drift with the relatively rapid speed of evolution in some cave animals (Niemiller et al., 2008; Klaus et al., 2013; Zwang and Li, 2013; Fumey et al., 2018; Herman et al., 2018). Accordingly, it has also been postulated that pigmentation loss may be adaptive either based on positive selection for energy conservation or indirect selection dependent on pleiotropic functions of pigmentation. In cave habitats that largely lack primary production and exhibit scarce food resources, selection will favor individuals lacking pigmentation as a means of reducing the overall energy expenditure. The hypothesis of energetic saving is a common explanation for trait regression in cave dwellers (Barr, 1968, Jeffery, 2009, Gross et al., 2016, Culver et al., 2023), but requires experimental support.

Numerous pleiotropic functions of melanogenesis have been discovered, including behaviors, thermoregulation, life history and immunity (reviewed in True, 2003, Ducrest et al., 2008, Wittkopp & Beldade 2009), and indirect selection for albinism via pleiotropy has been suggested in *Astyanax* cavefish (Bilandžija et al., 2013a; Bilandžija et al. 2018, Jeffery et al., 2016; O’Gorman et al., 2021). The mutated *oca2* gene (Protas et al., 2006) which controls albinism by functioning in L-tyrosine processing in the melanosome during the first step in melanin biosynthesis (Manga et al., 2001; Chen et al., 2002) has pleiotropic effects on CAT levels and related physiology and behaviors and is under positive selection in three cavefish populations (Bilandžija et al., 2013a; Bilandžija et al. 2018, Jeffery et al., 2016; O’Gorman et al., 2021). Additionally, trade-offs between melanogenesis and metabolic adaptations to a low nutrient environment have also been suggested in *A. mexicanus* cavefish via cis-regulatory changes in a gene involved in tyrosine metabolism (Krishnan et al., 2022). The adaptive hypothesis for the evolution of albinism requires targets in which natural selection can act to increase fitness in cave adapted species, Thus, another objective of this study was to identify potential benefits of albinism in cave animals.

Our results show that albinism has evolved convergently by disruption of the first step in melanin synthesis in diverse cave animals ranging from annelids and mollusks to vertebrates. Remarkably, despite the loss of melanin in albino cave animals, cells capable of melanin synthesis are conserved in patterns resembling their close surface relatives. Furthermore, in one cave-adapted vertebrate and two cave-adapted invertebrate species, cells capable of synthesizing melanin are recruited to the sites of injuries, suggesting important roles in innate immunity. In addition, we provide the first experimental evidence supporting the hypothesis that the loss of pigmentation could represent an energy saving adaptation. We show that melanin synthesis has a marked cost in the teleost *Astyanax mexicanus*, a single species consisting of pigmented surface fish and multiple populations of depigmented and albino cavefish (Jeffery, 2020). Lastly, we find that the disruption of melanin synthesis at its first step enhances dopamine in multiple depigmented cavefish populations, supporting a widespread tradeoff between the melanin and CAT synthesis pathways in *A. mexicanus*. The consistent block at the first step of melanin synthesis across cave-dwellers from different phyla is not compatible with the random process of neutral mutation and genetic drift. Furthermore, the identification of multiple advantages for the loss of pigmentation support the role of natural selection in the convergent evolution of albinism in cave animals.

## Materials and Methods

### Biological materials

*A. mexicanus* surface fish and cavefish were obtained from the Jeffery Laboratory Colony at the University of Maryland, College Park. Cavefish populations used in this study were Pachón, Tinaja, Los Sabinos, and Molino. Fertilized eggs and larvae were obtained by spawning as described by Ma et al. (2021a). F1 hybrids were produced by crossing surface fish X Pachón cavefish. The albino eyed (AE) strain was prepared by artificial selection in a four generation surface fish x Pachón cavefish intercross (Ma et al., 2021b). Methods for fish handling and experimental procedures were approved by the University of Maryland IACUC and conformed to USA National Institutes of Health guidelines.

The polychaetes *Marifugia cavatica* and *Ficopomatus enigmaticus* were collected in Tounjčica Cave, Tounj, Croatia and in the Krka River estuary, Skradin, Croatia, respectively. The mollusks *Congeria kusceri*, *Dreissena polymorpha*, and *Melledela werneri* were collected in Pukotina u tunelu polje jezero – Peračko Blato, Peračko Blato, Croatia; Jarun Lake, Zagreb, Croatia and Ostaševica Cave, Mljet, Croatia, respectfully. The crustaceans *Asellus aquaticus* and *Gammarus sp.* were collected in the Sava River, Zagreb, Croatia. The planarian *Polycelis sp*. was collected in Majerovo vrilo, Otočac, Croatia, and an uncharacterized species of cixiid planthopper was collected at Sedrena špilja, Oklaj, Croatia. The fin clip of amphibian *Proteus anguinus* was collected in Oko Cave, Bosnia and Herzegovina. Permits for these collections were UP/I-612-07/20-48/83, 517-05-1-1-20-5 (Ministry of Environmental Protection and Energy); UP/I-352-04/22-08/96, 517-10-1-1-22-5 (Ministry of Economy and Sustainable Development); from the Department for Nature Protection of the Republic of Croatia, 04-23-880/18 (Federal Ministry of Environment and Tourism, Federation of Bosnia and Herzegovina). *Ambylopsis spelaea* and *Typhlichthys subterraneus* fin clips were collected under scientific permits issued by the states of TN (no. 1605) and KY (no. SC1211135), USA and kindly provided by Prof. Daphne Soares and Matthew L. Niemiller.

### Pigment identification assays

To determine whether albinism in our target species was a result of disruption of melanogenesis and not some other pigment type, we developed simple pigment solvent assays. Specimens were fixed in 5% formalin for 1 h, rinsed in PBS, and subsequently incubated in solvents, and the effects on dark pigmentation were determined by microscopy. We used acetone, which extracts carotene (Ghidalia, 1985), formic acid, which dissolves ommochome and porphyrin but not melanin pigment (Ghidalia, 1985; Needham, 1974), and the melanin bleach kit (Sigma, St. Louis, USA) to extract melanin, but not carotene or ommochrome pigment, in our test species. As controls, we incubated tailfin clips of *A. mexicanus* surface fish, which have xanthophores containing carotene pigment and melanophores containing melanin pigment. Additionally, we used *A. aquaticus* with ommochrome pigmentation, *G. minus*, which produces melanin at wound sites, cixiid insects with melanin pigmentation, and the planarian (*Polycelis sp*.) with porphyrin pigmentation in the body and melanin pigment in the eyes.

### Melanogenic substrate assay

To identify the step of melanin synthesis that was disrupted in albino cave animals we performed the melanogenic substrate assay, as described in Bilandžija et al (2012). Albino cave species and closely related pigmented surface species were lightly fixed in 5% formalin in PBS for 1 hr. After several cycles of washing in PBS, they were incubated in a 0.1 % buffered solution of L-DOPA or L-tyrosine. Some of the fixed L-DOPA positive albino cave species and closely related surface species were embedded in Paraplast and sectioned at 10 µm using a Microtome. Whole specimens were viewed using a Zeiss Axioskop compound microscope.

### Wounding procedures

In *A. mexicanus* surface fish and Pachón cavefish larvae wounds were inflicted in the dorso-lateral musculature just posterior to the yolk sac at 5-10 days post fertilization (dpf) using sharp microcapillaries. In adults, wounds were produced by severing a distal part of the tail fin. The wound sites of surface fish were observed under a stereomicroscope for several days. The cavefish were observed by stereomicroscopy and then fixed for 1 hour at 4°C with 5% formalin in PBS at several consecutive time points during wound healing. Following fixation, the tissue was subjected to the melanin substrate assay as described above, and the L-DOPA rescued cells were observed by stereomicroscopy. Macrophage distribution around the wound sites of surface fish and cavefish was followed by neutral red staining as described previously (Ma et al., 2020b). The polychaetes *F. enigmaticus* and *M. cavatica* and the bivalves *D. polymorpha* and *C. kusceri* were wounded by making a lesion in different body parts using microcapillaries. The wound sites of *F. enigmaticus* were observed under a stereomicroscope for 24-72 hours. The wound sites of *M. cavatica, D. polymorpha, and C. kusceri* were observed at 24 and 48 hours after wounding, fixed for 1 hour at 4°C with 5% formalin, and subjected to the melanogenic substate assay.

### Morpholino knockdown of the *oca2* gene

The *A. mexicanus oca2* gene was knocked down by injecting a translation-blocking morpholino (MO) (5’-CTTGTTCTCCAAATACATCACACCT-3’) corresponding to bp −7 to +18 of the gene, as described by Bilandžija et al. (2013a, 2018). The *oca2* MO and a control MO (5’-CCTCTTACCTCAGTTACAATTTATA-3’) were designed and provided by Gene Tools (Summerton, OR, USA). We injected 400 pg of each MO into 1-2 cell embryos, which provided maximal inhibition of melanophore development with minimal effects on morphology.

### Oxygen consumption measurements

We tested whether melanin synthesis requires energy in *A. mexicanus* by comparing oxygen consumption in *oca2* and control morphant larvae. The morphant larvae were placed in air-tight vials filled with culture system water and incubated at room temperature for about 30 hours. The amount of oxygen left in vials was measured using a Membrane Inlet Mass Spectrometer (Bay Instruments, Easton, MD, USA). Vials without larvae were used as blanks.

The value of oxygen consumption of each larva was calculated by subtracting the amount of oxygen measured in its vial from the mean of oxygen level of 5 blank vials. Because MO knockdown is a transient method, some pigmentation eventually breaks through the block. Therefore, following the oxygen measurements, individual *oca2* morphant larvae were inspected for the presence or absence of pigmentation and separated into *oca2* albino and *oca2* pigmented morphant groups.

### High-performance liquid chromatography

Whole brains were dissected from adult fish and collected for high performance liquid chromatography (HPLC). Dopamine (DA) and 3,4 dihydroxyphenylacetic acid (DOPAC) were analyzed using HPLC with electrochemical detection as described previously (Renner and Luine, 1986), with minor modifications. Brains were placed into 100 µL of acetate buffer (pH 5.0) containing the internal standard a-methyl DA (Merck & Co., Kenilworth, NJ), sonicated and stored at-80 °C. Prior to analysis, 4 µL ascorbate oxidase (1 mg/mL, Sigma-Aldridge) was added to the samples followed by centrifugation at 17 000 x g for 15 min. A Waters Alliance e2695 was used to inject 50 µl of the supernatant onto a C18 4 mm NOVA-PAK column (Waters Associates, Milford, MA) held at 30 °C. Electrochemical detection was accomplished using an LC 4 potentiostat and glassy carbon electrode (Bioanalytical Systems, West Lafayette, IN) set at 0.5 nA/V with an applied potential of 0.7 V versus an Ag/AgCl reference electrode. The pellet was solubilized in 400 µL of 0.4 N NaOH and protein content was analysed according to Bradford [42]. ACSW32 data program (DataApex, Czech Republic) was used to determine monoamine concentrations in the internal standard mode using peak heights calculated from standards. DA and DOPAC concentrations were corrected for injection versus preparation volumes and divided by micrograms protein to yield picogram monoamine per microgram protein. Data was tested for differences using a One Way Analysis of Variance because it passed normality and equal variance tests (SIGMASTAT version 3.5, Systat Software, San Jose, CA). In analyses that revealed a significant effect between groups, Dunn’s method was used to conduct pairwise comparisons. Application of Grubb’s test [43] to detect outliers resulted in the deletion of data from one brain from the dataset. Significance levels for all statistical tests were set at p = 0.05.

## Results

### Melanin pigmentation in surface relatives of albino cave species

Dark pigment in vertebrates is based on melanin, but invertebrates use melanin and/or other pigment types for dark coloration, most commonly ommochromes and porphyrins (Bandaranayake, 2006). Therefore, to select cave species for this study, we first identified those with surface-dwelling relatives showing melanin pigmentation. Assays were developed in which surface species were extracted with solvents that dissolve different types of pigment: acetone for pigments soluble in organic solvents such as carotenes, formic acid for ommochromes and porphyrins, and a bleach treatment (melanin bleach) for melanin. We tested the specificity of these solvents in *A. mexicanus* surface fish, which contain brown to black melanin-containing melanophores and orange carotene-containing xanthophores (Fig. 1A) (Jeffery et al., 2016). Acetone removed xanthophores, but not melanophores (Fig. 1B), identifying carotene as the xanthophore pigment, formic acid did not affect xanthophores or melanophores (Fig. 1C), consistent with the absence of ommochromes and porphyrins, and melanin bleach extracted melanophores but not xanthophores (Fig. 1D). These results confirm melanin as the dark pigment in *A. mexicanus* and illustrate the specificity of the pigment solvent assays.

**Figure 1.**
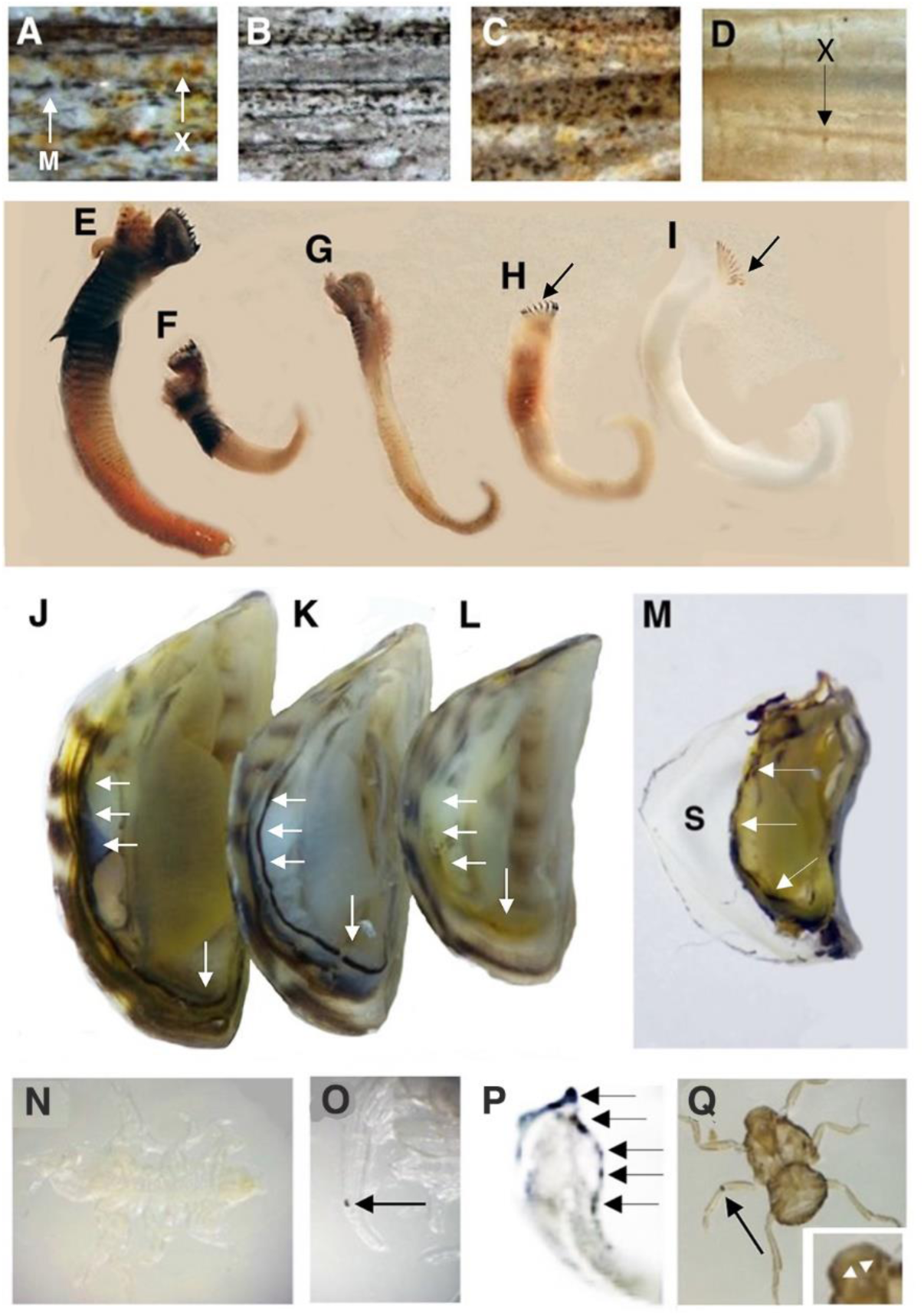
Pigment solvent assays distinguish between dark pigmentation types in vertebrate and invertebrate species. A-D. *Astyanax mexicanus* surface fish tailfins. A. Untreated control showing melanophores (M) and xanthophores (X). B. Acetone removes xanthophores. C. Formic acid has no effects on melanophores or xanthophores. D. Melanin bleach removes melanophores but not xanthophores (X). E-I. *Ficopomatus enigmaticus.* E. Untreated control showing dark pigmentation in tentacles and segments. F, G. Acetone (F) or formic acid (G) do not extract dark pigmentation. H, I. Melanin bleach decreases (30 min treatment, H) or removes (12 h treatment, I) dark body pigmentation but not tentacle pigmentation (arrows). J-M. *Dreissena polymorpha.* J. Untreated control showing dark pigmentation in the shell and body, including a pigment band along the edge of the mantle (arrows). K. Acetone does not affect dark shell pigmentation or mantle pigmentation (arrows). L. Melanin bleach removes the mantle band pigmentation (arrows) but not shell pigmentation. M. Formic acid dissolves the shell (S) but does not affect dark mantle pigmentation (arrows). N. *Asellus aquaticus*: formic acid removes ommochrome body pigmentation. O. *Gammarus minus:* formic acid removes body pigment but not melanin in a wound on the antenna (arrow). P. *Polycelis sp.*: formic acid removes body pigmentation but not dark pigment in multiple eyes (arrows). Q. Unknown cixiid insect species: formic acid does not remove body pigment, eye pigment (inset), or a melanized appendage wound (arrow).

As additional controls, the solvent assays were used in surface invertebrate species known to contain ommochrome or porphyrin pigmentation in addition to melanin. In the isopod *Asellus aquaticus*, which contains only ommochrome pigmentation (Needham and Brunet, 1957), dark body coloration was completely removed by formic acid (Fig. 1N). In arthropods, melanin is synthesized at wound sites as a response to injury (Bilandžija et al., 2017). Thus, in the amphipod *Gammarus minus* body pigment, but not melanized wound pigment, was removed with formic acid (Fig. 1O). In the surface planarian *Polycelis sp*., porphyrin body pigmentation (Needham, 1965) was extracted with formic acid (Fig. 1P), but dark pigmentation in the multiple eyes was not affected (Fig. 1Q). This is consistent with planarian eyes containing melanin (Hase et al, 2006). Lastly, in a surface-dwelling cixiid insect species, formic acid did not extract body or wound pigment, suggesting a melanin basis for dark pigmentation (Fig. 1Q and inset). However, eye pigment was extracted suggesting it is likely ommochrome, a common eye pigment in arthropods (Osanai-Futahashi et al 2012., Dontsov &. Ostrovsky 2023). These results show that pigment solvent assays can distinguish between melanin and other dark pigments in the same animals.

Using the assays developed above, we next surveyed pigment types in different surface-dwelling invertebrate species with dark pigmentation. We found that dark pigmentation in body segments of the polychaete annelid *Ficopomatus enigmaticus*, a close surface relative of cave-dwelling *Marifugia cavatica* (Kupriyanova et al., 2009), was extracted with melanin bleach, but not formic acid or acetone, indicative of melanin pigmentation (Fig. 1 E-I). In the surface bivalve mollusk *Dreissena polymorpha*, a close relative of the cave-dwelling *Congeria kusceri* (Bilandžija et al., 2013b), dark pigmentation at the edge of the mantle and gills was also extracted with melanin bleach, but not acetone or formic acid, confirming melanin pigmentation (Fig. 1 J-M). In contrast, dark coloration in *F. enigmaticus* tentacles (Fig. 1 H, I) and in the shell and other body parts of *D. polymorpha* (Fig. 1 L) was resistant to extraction with melanin beach, and thus not confirmed as melanin. In *D. polymorpha,* extraction of dark pigment stripes in the shell by formic acid (Fig. 1M) suggests that they are composed of either ommochromes or porphyrins.

Together, these results underscore the necessity of testing dark pigmented species for the presence of melanin before related albino cave species are concluded to be deficient in melanin synthesis. We conclude that in addition to vertebrates, melanin is a body pigment in some annelid and mollusk species, permitting them to be coupled with closely related albino cave species in the pigmentation studies described below.

### The first step of melanin synthesis is disrupted in diverse albino cave species

It was previously shown that melanin biosynthesis is blocked at the first step of the pathway, the conversion of L-tyrosine to L-DOPA, in the albino Pachón form of *A. mexicanus* cavefish (McCauley et al., 2004) and in two albino cave cixiid species (Bilandžija et al., 2012), suggesting that disruption at this step could occur in diverse cave animals. To address this possibility, the melanogenic substrate assay (McCauley et al., 2004; Bilandžija et al., 2012), in which albino species are challenged to produce melanin when supplied with exogenous L-tyrosine or L-DOPA substrates, was applied to additional albino cave animals whose surface relatives contain melanin or are embedded in surface lineages already known to have melanin-based pigmentation (teleosts and amphibians). Treatment with L-DOPA, the second substrate in the melanin pigment pathway, restored pigmentation in the cave mollusks *Melledela werneri* (Fig. 2B) and *C. kusceri* (Fig. 2D), the cave polychaete *M. cavatica* (Fig. 2F), the cavefishes *Typhlichthys subterraneus* (Fig. 2H) and *Amblyopsis spelaea* (Fig. 2J), and the cave salamander *Proteus anguinus* (Fig. 2L). In contrast, the provision of exogenous L-tyrosine, the first substrate in the melanin pigment pathway, did not restore melanin biosynthesis in any of these albino cave species (Fig 2A, C, E, G, I, K). Restoration of pigmentation by L-DOPA, but not L-tyrosine, indicates a block in melanin synthesis at the first step of the pathway in diverse albino cave species and that all the downstream steps are intact and functional.

**Figure 2.**
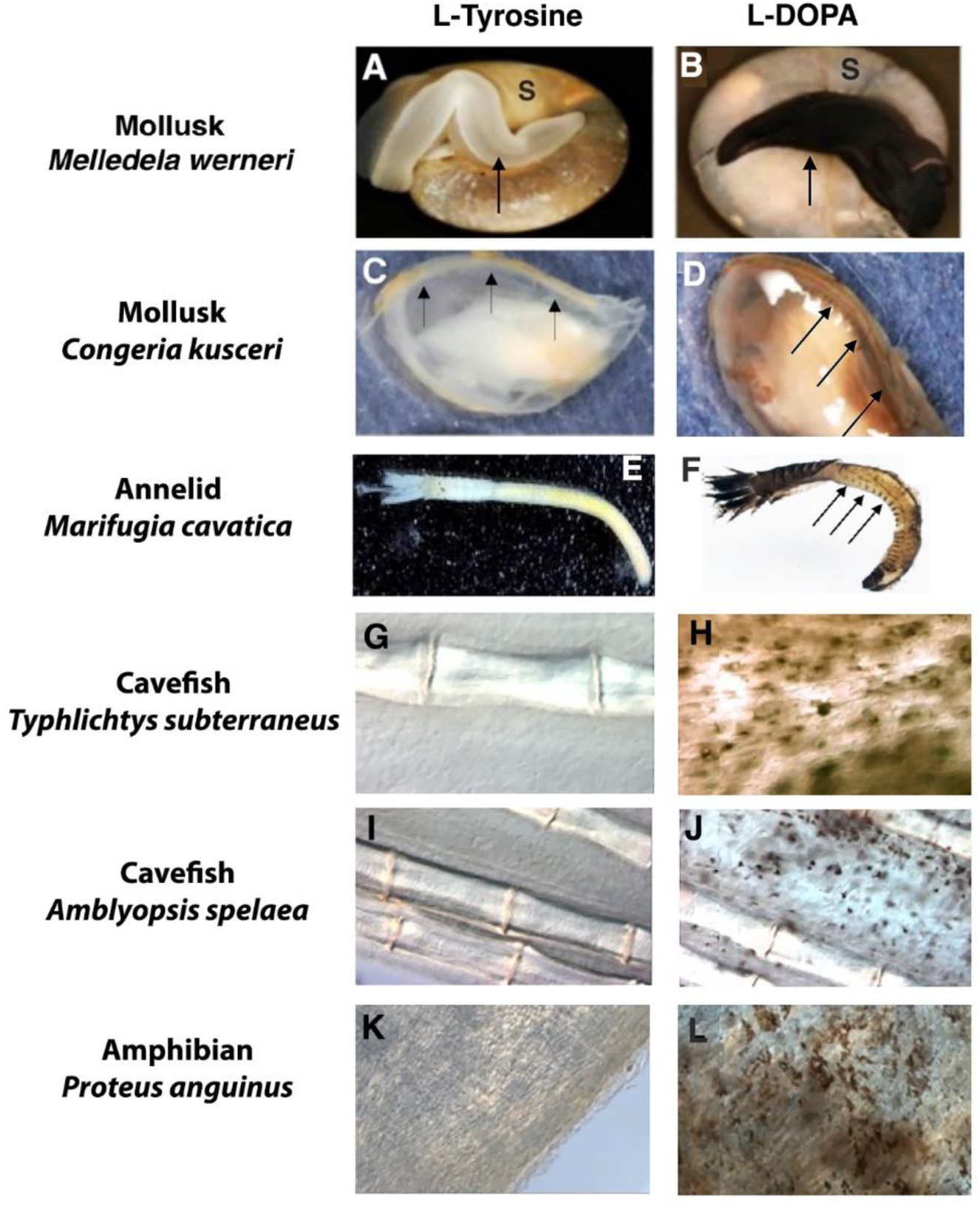
Diverse albino cave-adapted species are blocked at the first step of melanin synthesis. A, C, E, G, I, K. The addition of L-tyrosine, the first substrate in the melanin synthesis pathway, does not rescue melanin pigmentation in (A) the albino cave gastropod *Meledella werner,* (C) the albino cave bivalve *Congeria kusceri,* (E) the albino cave polychaete *Marifugia cavatica*, (G, I) the albino cavefish (G) *Ambylopsis spelaea* and (I) *Typhlichthys subterraneus,* and (K) the albino cave salamander *Proteus anguinus.* B, D, F, H, J, L. The addition of L-DOPA, the second substrate in the melanin synthesis pathway, rescued melanin pigmentation in each albino cave species. S in A, B: shell. Arrows in A, B: Protruding foot. Arrows in C, D: rescued pigmentation along the edges of the mantle. Arrows in F: rescued pigmentation stripes in the segmented posterior body

### The pattern of restored melanin pigmentation in cave species resembles natural dark pigmentation in surface relatives

To determine the patterns of pigment restoration in albino cave species, the distribution of L-DOPA rescued pigment cells in albino cave animals was compared to the natural pigmentation patterns in their surface relatives. In the cave polychaete *M. cavatica* (Fig. 3A), L-DOPA rescued pigmentation in a segmental pattern resembling the natural pattern of body pigmentation in surface *F. enigmaticus* (Fig. 3B). In the cave bivalve *C. kusceri*, L-DOPA rescued melanin in cells located at the edge of the mantle and gill where melanin pigment is normally located in surface *D. polymorpha* (Fig. 3C, D). In *A. mexicanus* Pachón cavefish larvae (Fig. 3E) and adult tailfins (Fig. 3F), L-DOPA rescued pigment cells were distributed similarly to melanophores in surface fish larvae (Fig. 3G) and tailfins (Fig. 3H). Thus, despite the loss of melanin pigmentation, cells capable of melanin synthesis are conserved in these diverse albino cave animals and distributed in patterns resembling melanin pigment cells in closely related surface forms. These results suggest that the loss of pigmentation is due to disruption of the capacity for melanin synthesis and not the loss of cells originally responsible for producing pigmentation.

**Figure 3.**
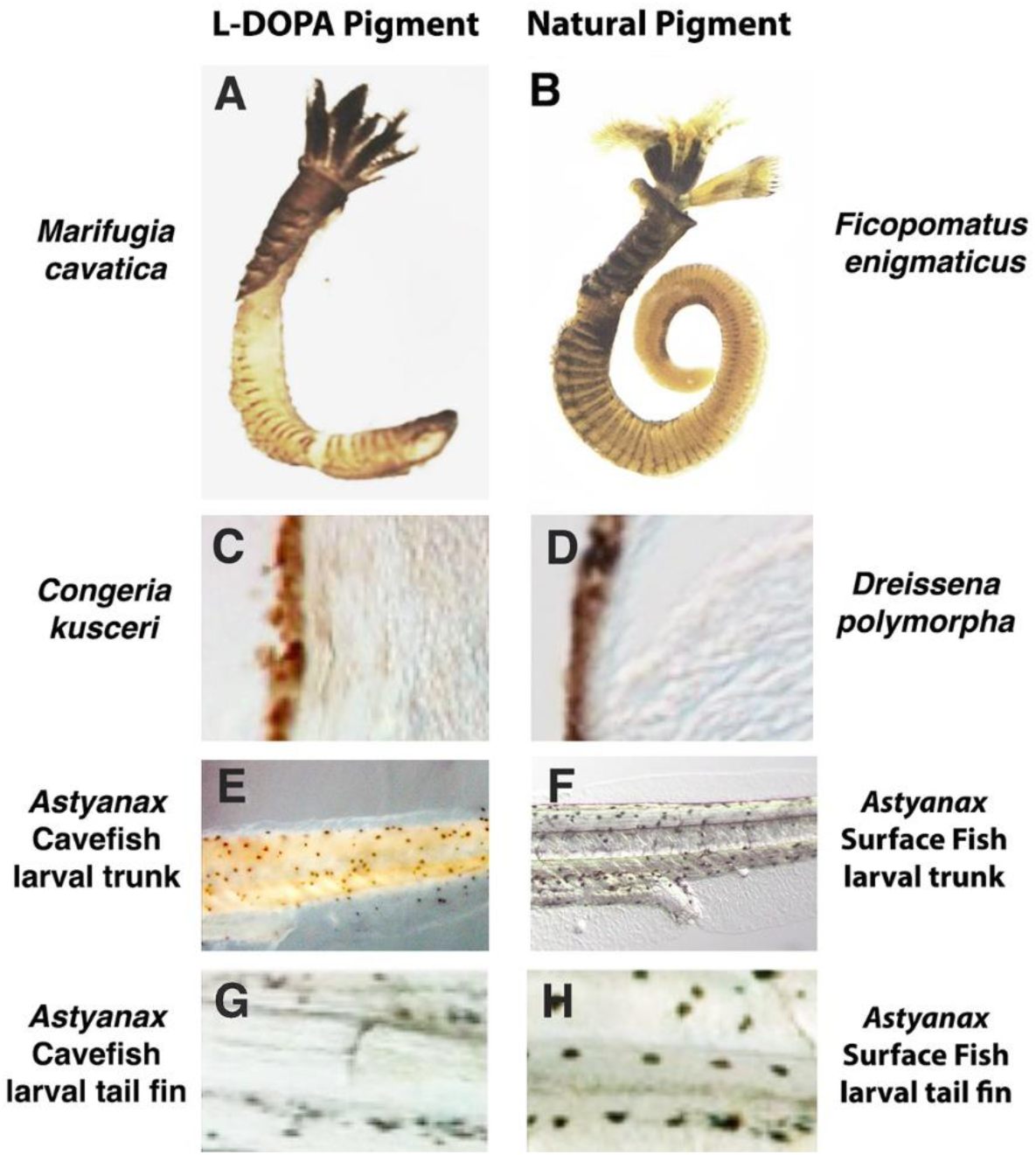
The pattern of L-DOPA-rescued melanin pigmentation in albino cave animals resembles natural melanin pigmentation in closely-related surface relatives. A, B. L-DOPA rescued body pigmentation in albino cave *M. cavatica* (A) and natural body pigmentation in surface *F. enigmaticus* (B). C, D. L-DOPA rescued pigment cells at the edge of the mantle in albino cave *C. kusceri* (A) and natural pigmentation at the edge of the mantle in surface *D. polymorpha* (B). E-H. *A. mexicanus* Pachón cavefish larval body (E) and tailfin (F) with L-DOPA-rescued pigment cells in surface fish larval body (G) and tail fin (H) with melanophores.

### Cells capable of synthesizing melanin pigment may function in innate immunity in albino cave animals

The conservation of cells capable of synthesizing melanin suggests that they have indispensable roles in albino cave species. In arthropods and other invertebrates, melanin is an important mediator of innate immunity (Sugumaran, 2002; Gimaldi et al., 2012; Mackintosh 2001, Christensen et al., 2005; Bilandžija, et al., 2017), but this function is not well known in vertebrates, which in addition to innate immunity have evolved an adaptive immune response. To address whether the cells capable of synthesizing melanin in albino cave species function in innate immunity, we inflicted injuries to surface and cave *A. mexicanus*, surface *F. enigmaticus* and cave *M. cavatica,* and surface *D. polymorpha* and cave *C. kusceri*, and then followed the appearance of melanin-containing pigment cells and L-DOPA-rescued pigment cells, respectively (Fig. 4).

**Figure 4.**
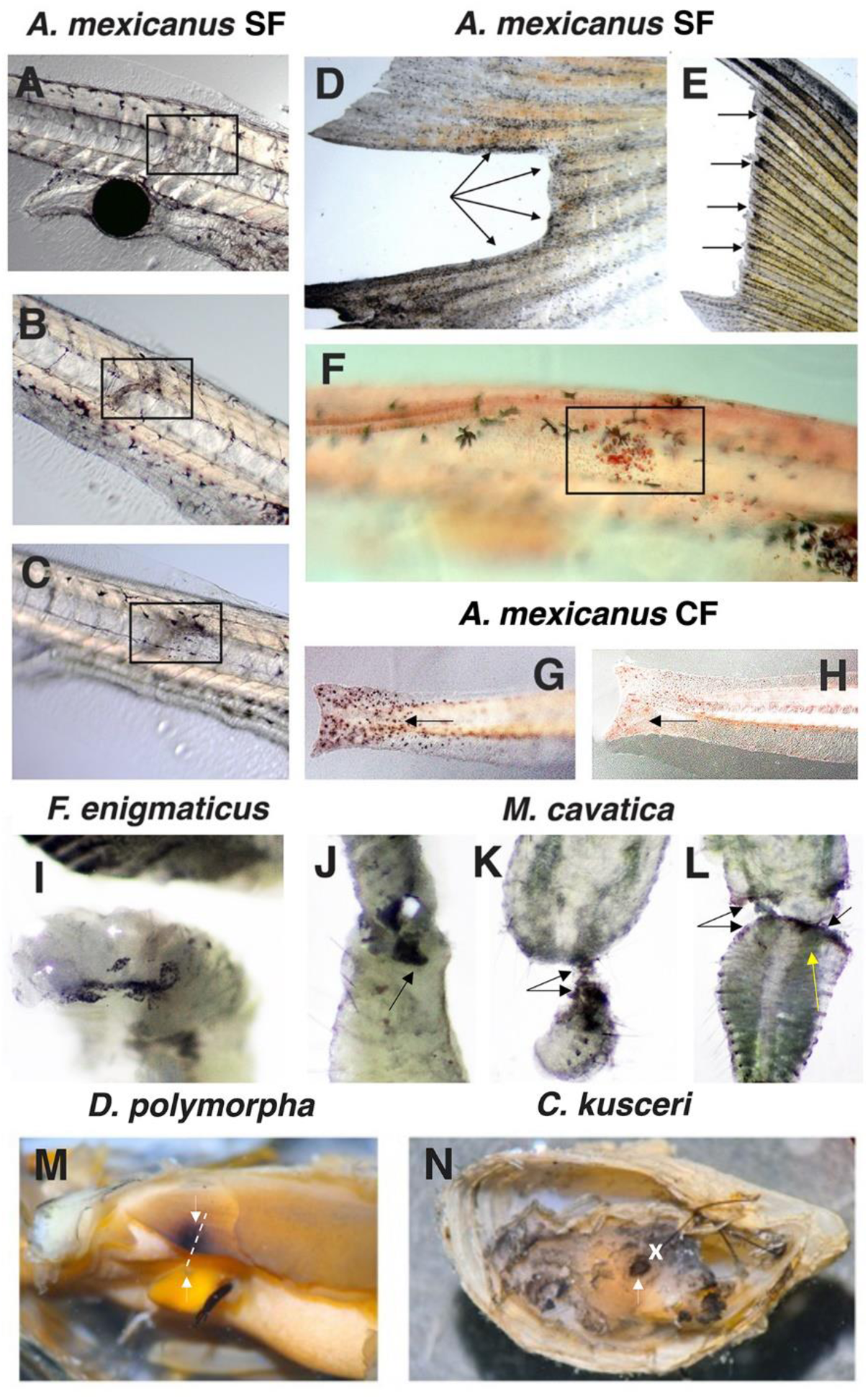
Melanin pigment and L-DOPA rescued cells are involved in injury repair in diverse cave albino species. A-C. The accumulation of melanophores at wound sites (boxes) in a 7-day post-fertilization (dpf) *A. mexicanus* surface fish larva shortly after injury (A), at 24 hrs. post injury (B), and at 48 hrs. post injury (C). D, E. L-DOPA-rescued pigment cells or melanophores appear at the edge of wounds (arrows) in adult *A. mexicanus* Pachón cavefish (D) and surface fish (E), respectively. F. Melanophores and neutral-red stained macrophages appear at a wound site (boxed area) in 7 dpf surface fish. G, H. L-DOPA-rescued pigment cells (G) and neutral red-stained macrophages (H) at the severed edges of tailfin wounds in 7 dpf Pachón cavefish. I-L. Black pigment cells in surface *F. enigmaticus* (I, arrow) and L-DOPA rescued pigment cells in cave *M. cavatica* (J-L, arrows) are present about 24 hr. after being wounded. Yellow arrow in L shows L-DOPA rescued cells in a segmented band. M, N. L-DOPA rescued pigmentation in *D. polymorpha* and *C. kusceri* (white arrows). Dashed line in M and X in N: wound sites.

Superficial wounds were inflicted in the trunks of 7-day post-fertilization (dpf) *A. mexicanus* larvae and adult tailfins and the subsequent distribution of melanophores in surface fish and L-DOPA-rescued cells in Pachón cavefish were followed by microscopy (Fig. 4A-H). In surface fish, melanophores accumulated at wound sites in both larvae and adults (Fig. 4A-C, E, F). L-DOPA-rescued pigment cells also appeared at wound sites in Pachón cavefish larvae and adults (Fig. 4 D, G, H). Notably, melanophores or L-DOPA rescued cells at the wound sites were joined by neutral red-stained macrophages (Fig. 4F), which serve as effectors in the innate immune system by eliminating damaged cells by programmed cell death (Hirayama et al., 2017). These results show that wounding mobilizes macrophages and melanin-containing or L-DOPA rescued pigment cells around the wound sites in surface fish and cavefish and suggest that the retention of cells with a capacity for melanin synthesis in cavefish may be explained by their function in innate immunity.

Wounds were inflicted along the segmented bodies of *F. enigmaticus* and *M. cavatica* (Fig. 4I-L) and in the mantles, gills and/or foot of *D. polymorpha* and *C. kusceri* (Fig. 4M, N) and subsequently examined by direct imaging or following L-DOPA rescue. In *F. enigmaticus*, black pigment cells appeared around the wounds, breaking the normal segmentation pattern of the melanin pigmented bands (Fig. 4I), and L-DOPA-rescued pigment cells were detected surrounding injuries in cave *M. cavatica* (Fig. 4J-L). Although black pigment cells were not detected around the wounds in *D. polymorpha,* in the latter and *C. kusceri,* L-DOPA rescued pigment cells did appear at wound sites (Fig. 4M, N). These results suggest that cells capable of melanin synthesis in albino cave annelid and mollusk species may be involved in injury repair.

In summary, our results suggest that cells capable of producing melanin may be conserved in diverse albino cave-adapted species because of important roles in innate immunity during tissue repair.

### Blocking melanin synthesis at its first step conserves energy in *A. mexicanus* embryos

Although energy conservation has been proposed as a driver of trait regression, including albinism in cave animals, this hypothesis has not been experimentally verified. Albinism in *A. mexicanus* Pachón cavefish is caused by a 130 base pair deletion in the *oca2* gene (Protas et al., 2006), a melanosome transmembrane protein, which may function in L-tyrosine transport into the melanosome or tyrosinase processing during melanin biosynthesis (Manga et al., 2001; Chen et al., 2002). As described previously (Bilandzija et al., 2013a), injection of an *oca2* morpholino (MO) into fertilized eggs results in the development of morphant larvae lacking or with substantially reduced pigmentation. Therefore, to explore the possible role of energy conservation in pigment loss, surface fish embryos were injected with *oca2* MO, the morphant larvae were separated into two groups, an albino group and a group with reduced pigmentation, and O_2_ consumption, as a measure of energetic/metabolic cost, was compared between the two *oca2* morphant groups and normally pigmented larvae injected with a control MO (Fig. 5). Albino *oca2* morphants consumed less oxygen than pigmented control morphants (Fig 5). Furthermore, *oca2* morphants with reduced pigmentation showed O_2_ consumption levels intermediate between albino *oca2* morphants and fully pigmented controls (Fig. 5). The results indicate that *oca2* knockdown and pigment loss substantially lowered the amount of energy expended during development of *A. mexicanus* surface fish. Therefore, the disruption of melanin synthesis at its first step could be an advantage for survival and a target of natural selection in the cave environment.

**Figure 5.**
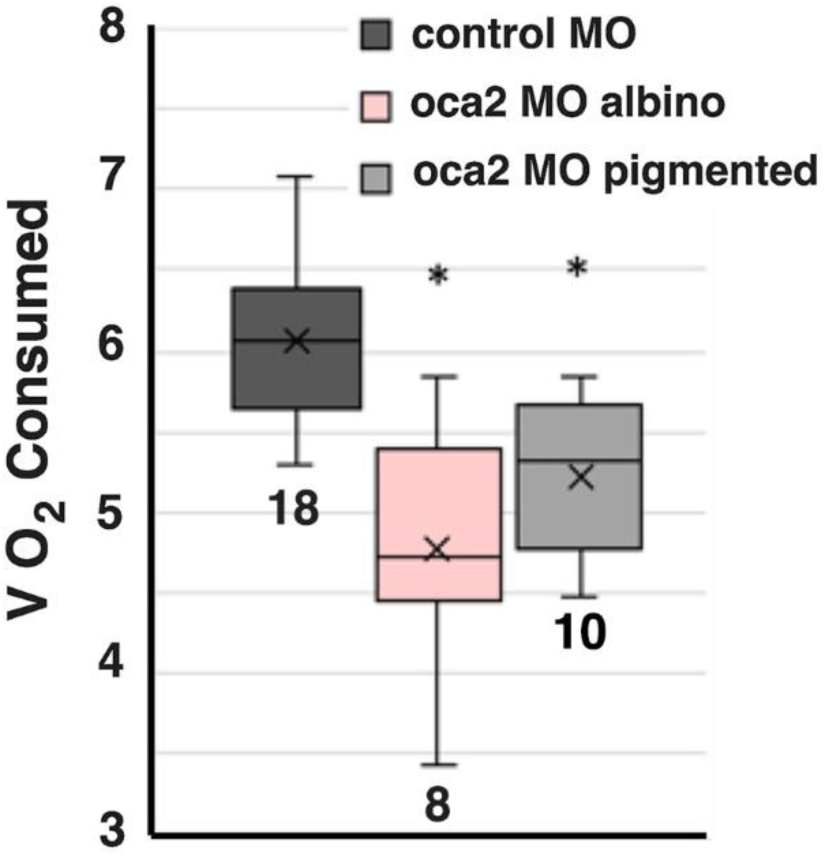
The effects of *oca2* knockdown on oxygen consumption in *A. mexicanus* larvae. Box and whisker plots showing 4-day post-fertilization larvae injected with *oca2* morpholino separated into albino (albino oca2 MO) and pigmented (pigmented oca2 MO) groups and compared with control morpholino (pigmented control M). The number of samples is shown at the bottom of each box. Asterisks represent significance at p < 0,001 according to One-way ANOVA with Bonferroni corrections. Boxes represent data values in between the first and the third quartile, whiskers represent minimum and maximum values, while the x represents the mean, and horizontal bar represents median.

### Melanin pigment reduction and albinism increases dopamine synthesis in *A. mexicanus*

It was previously shown that L-tyrosine and dopamine (DA) are increased in whole larvae and adult brains of Pachón cavefish compared to surface fish (Bilandžija et al., 2013a). Furthermore, silencing of *oca2* in the surface *A. mexicanus* caused an increase of L-tyrosine and DA levels suggesting a tradeoff between the melanin and CAT synthesis pathways (Bilandžija et al., 2013a). To further address this tradeoff, we compared DA levels in various lineages of *A. mexicanus*, including cave populations with various degrees of depigmentation and independent loss of pigmentation as well as hybrids resulting from artificial selection for albinism (Fig. 6).

**Figure 6.**
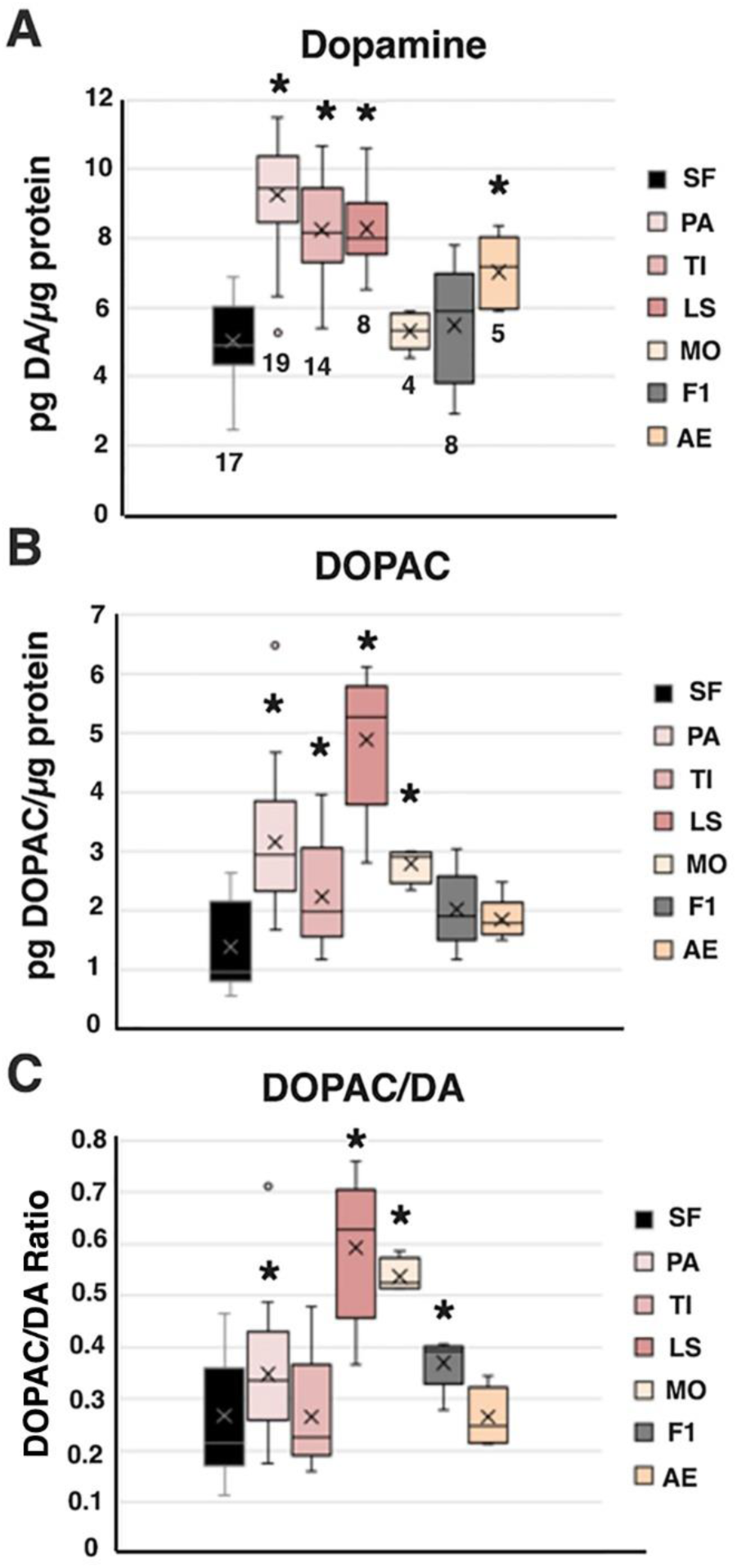
Dopamine levels and dopaminergic activity in the brains of different *A. mexicanus* cavefish populations and hybrids. A-C. Box-whisker plots showing dopamine (DA) (A), DOPAC (B), and dopaminergic activity (DOAC/DA) (C) in control surface fish (SF), Pachón, Tinaja, Los Sabinos, and Molino cavefish, surface fish X Pachón cavefish F1 hybrids, and, and the albino eyed strain (see figure legends on right). Boxes represent data values in between the first and the third quartile, whiskers represent minimum and maximum values, while the x represents the mean, and horizontal bar represents median of the combined dataset based on two HPLC runs. Separate statistics for each run can be found in S1 Table.

We measured DA and DOPAC concentrations and calculated DOPAC/DA ratios, the latter as metabolic indices of dopaminergic activity, in two HPLC experiments. Several groups (SF, PA and TI) were included in both runs, and since there was no statistically significant difference in results between the two runs based on T-tests, we show combined results in the Figure (Fig. 6A-C) and statistics for each run separately in Supplement (S1 Table). In the first experiment, DA and DOPAC/DA were determined in brains isolated from the Tinaja, Los Sabinos, and Molino forms of *A. mexicanus* cavefish, which show different levels of melanin pigmentation (medium levels, low levels, and albinism, respectively), and compared to brains of albino Pachón cavefish, fully-pigmented surface fish, and pigmented surface fish x Pachón F1 hybrids. The results showed increased levels of DA in brains from depigmented Pachón, Tinaja, and Los Sabinos cavefish relative to pigmented surface fish and F1 hybrids. Although albino Molino cavefish brains showed no differences in DA levels (Fig. 6A), increased DOPAC/DA ratios suggested that dopaminergic activity is higher in the brains in Molino cavefish (Fig. 6B). In the second HPLC experiment, increased DA was confirmed in Tinaja and Pachón cavefish brains, and increased DA was also observed in brains of the *A. mexicanus* albino eyed (AE) strain, which was produced by crossing Pachón cavefish by surface fish followed by multiple rounds of artificial selection for albino progeny with fully developed eyes (Ma et al., 2020a). These results show that DA or dopaminergic metabolism is increased across multiple de-pigmented albino cavefish populations and artificially-selected albino hybrids, suggesting a widespread tradeoff between the melanin synthesis and CAT pathways in *A. mexicanus*.

## Discussion

This study shows that the convergent loss of pigmentation is controlled by the same mechanism in diverse cave animals, supporting the hypothesis that the evolution of albinism could be explained as an adaptation involving natural selection (Bilandžija et al., 2013a; 2018; O’Gorman, 2021) rather than as a stochastic mechanism involving neutral mutations and genetic drift (Juan et al., 2010; Wilkens and Strecker, 2017). The evidence stems from the broad convergence of a block at the first step of the melanin pathway and the discovery of multiple potential advantages in disrupting melanin biosynthesis at this step, which could serve as targets of natural selection in albino cave animals.

As primary biological subjects in this study, we chose the cave polycheate *M. cavatica*, the cave mollusks *C. kusceri* and *M. werneri*, the cave salamander *P. anguinus*, and three different cavefish species, including the classic cavefish model *A. mexicanus* (Jeffery, 2020). Based on assays developed to distinguish darkly pigmented species containing melanin from those containing porphyrins or ommochromes, we determined that the albino cave species covered in this study are closely related to surface dwelling species containing melanin pigment, and thus albinism in these species is a consequence of distrupting melanin synthesis. Accordingly, we employed the melanin substrate assay (McCauley et al., 2004), in which albino species are challenged to synthesize melanin following the provision of L-tyrosine, the first substrate, or L-DOPA, the second substrate in the pathway, to identify the step of melanin synthesis blockage. In each of the diverse cave species examined, we found that L-DOPA, but not L-tyrosine, rescued melanin synthesis. In addition, previous studies showed that *A. mexicanus* Pachón cavefish and two independently evolved cave cixiid species from Croatia and Hawaii, are also defective in the first step of the melanin synthesis pathway (McCauley et al, 2004; Bilandžija et al., 2012). Thus, melanin synthesis is disrupted at its first step in a wide range of cave-adapted animals. Since most cave adapted animals fit into one of the taxonomic groups covered in our studies, the results suggest that a block at the first step of the melanin synthesis pathway could be ubiquitous among albino cave animals.

According to the genetic drift hypothesis (Juan et al., 2010; Wilkens and Strecker, 2017), stochastic mutations would be expected to cause albinism by disrupting any step in the melanin synthesis pathway. However, this scenario was not supported by our results. Instead, only one step, the first step in the pathway, was always disrupted, supporting this single step as a frequent target of natural selection. Moreover, the ability to rescue pigmentation by the provision of L-DOPA substrate suggests that all the downstream steps and their effectors are present and functional. A potential caveat of this conclusion could be that the first step of the pathway might be the only step where disruption is possible. However, this constraint can be rejected by evidence showing that multiple steps in melanin synthesis can be targeted for mutation. For example, spontaneous or laboratory induced *Drosophila* mutations disrupt the melanin synthesis pathway at several early and late steps (Minakhina and Steward, 2006). Likewise, human albinism is caused by mutations in the early (OCA1 and OCA2 albinisms) or late (OCA3 and OCA4 albinisms) steps in melanin synthesis (Oetting and King, 1999). Thus, we conclude that blocking melanin synthesis at its first step, rather than at any other place in the pathway, may be a conserved feature of albino cave animals.

Our discovery of cells with the capacity for melanin synthesis after L-DOPA provision in divergent albino cave animals, and patterns representing their likely ancestral distributions as evident in closely related surface species, suggest important functions other than body pigmentation. To explain these results, we tested the hypothesis that melanin pigment cells have anti-microbial properties (Mackintosh, 2001), which is supported by melanophore recruitment to the sites of tissue damage in zebrafish (Lévesque et al., 2013). This hypothesis has not been addressed previously in albino cave animals. When *A. mexicanus* surface fish and cavefish larval bodies or adult tailfins were wounded, melanophores or cells capable of melanin synthesis after supplying L-DOPA substrate, respectively, were recruited to the sites of injury along with macrophages, which are known to be attracted to damaged tissues as part of the innate immune response (Hirayama et al., 2017). L-DOPA rescued pigment cells were also detected at wounds in cave *M. cavatica* and cave *C. kusceri*, suggesting that despite their albino phenotypes, cells with a cryptic capacity for melanin synthesis may be involved in innate immunity during wound healing in albino cave invertebrates.

Energetic savings is another hypothesis that has been proposed to explain the regressive evolution of cave animals (Barr, 1968, Jeffery, 2009, Gross et al. 2016, Culver et al. 2023). This hypothesis has received experimental confirmation only in the case of eye regression, where eye loss in *A. mexicanus* has been shown to reduce the overall metabolic expenditure (Moran et al., 2015). However, it has not been tested for pigmentation loss. If there is a cost for melanin synthesis, then it would be beneficial to block the melanin synthesis pathway early or before it is initiated. Accordingly, a block would be most effective in conserving energy at the first step of the pathway, for example by preventing the transport of L-tyrosine into the melanosome, which is a potential role of the OCA2 protein in vertebrates (Toyofuku et al., 2002). Although energy conservation is an appealing explanation for blocking melanin synthesis at the first step, until now this hypothesis also had not been directly tested in any cave animal. Our studies, in which oxygen consumption as a measure for energetic cost was measured in albino and depigmented *A. mexicanus oca2* morphants, show that melanin synthesis has a metabolic cost, at least in this system, and provides the first experimental evidence supporting the role of energy conservation in the regression of pigmentation in a cave animal. Together with studies showing that *oca2* CRISPR mutations may confer a feeding advantage in the dark cave environment (Choy et al., 2025), our studies involving energy conservation support the adaptive hypothesis for the evolution of albinism in *Astyanax* cavefish. We were unable to extend these studies to invertebrate cave species because the key gene(s) mutated to disrupt melanin synthesis has been identified only in *A. mexicanus*.

Altered behaviors and physiological functions in *A. mexicanus* o*ca2* morphants and CRISPR mutants may be dependent on CAT functions in the brain and other parts of the body (Bilandžija et al., 2013a; 2018, O’Gorman et al., 2021). Because the CAT and melanin synthesis pathways begin with the same substrate, L-tyrosine, a tradeoff has been proposed in which surplus L-tyrosine available from blocking melanin synthesis at its first step is used to increase CAT synthesis (Bilandžija et al., 2013a), thus providing a beneficial trait that could be a target of natural selection. Accordingly, it has been shown that knockdown of *oca2* increases L-tyrosine and DA levels in *A. mexicanus* surface fish larvae and that naturally elevated levels of DA are present in the brains of Pachón albino cavefish (Bilandžija et al., 2013a). The adaptive value of increased DA in cave evolution of *Astyanax* is not entirely clear. However, we reasoned that, if adaptive, increased DA would evolve in multiple independent cavefish lineages (Bilandžija et al., 2020), similar to higher starvation resistance and triglyceride content (Aspiras et al., 2015). Indeed, we found that increased DA or dopaminergic activity is present in four different cavefish populations. Our results expand and are consistent with a previous study of the melanin-CAT tradeoff in *A. mexicanus* (Bilandžija et al., 2013a) by documenting increases in DA levels and/or dopaminergic activity in the brains of additional depigmented and albino cavefish populations and albino hybrids produced through artificial selection. Therefore, we conclude that another advantage of blocking melanin synthesis at its first step, at least in *A. mexicanus*, is the enhancement of DA levels, which could be used to bolster or alter behavioral responses for better fitness in the cave environment.

In summary, we have discovered multiple advantages for blocking melanin synthesis at its first step. Some of these advantages could be widespread in albino cave animals, whereas others could be important in only some cave adapted species. Nevertheless, these results support an adaptive hypothesis based on natural selection for the loss of pigmentation in cave animals. The adaptive hypothesis is consistent with the rapid evolution of *A. mexicanus* cavefish, which may have diverged from surface fish only about 30,000 years ago (Fumey et al., 2018; Herman et al., 2018), and the rapid evolution of specialized cave traits in diverse cave-adapted animals (Niemiller et al., 2008; Klaus et al., 2013; Zwang and Li, 2013). The adaptive hypothesis is also supported by historical field studies in Pachón Cave (Borowsky, 2023). In the 1960s, researchers encountered *Astyanax* fish in Pachón Cave ranging from fully dark to albino individuals with all the different intermediate levels of pigmentation, suggesting that an episode(s) of introgression had occurred with nearby populations of surface fish. This is not an isolated case of mixing and hybridization between cavefish and surface fish as introgression frequently occurs in other caves harboring *A. mexicanus* (Mitchell et al., 1977). However, by the 2000s, all the fish observed in Pachón Cave were found to be albino (Borowsky, 2023). Genetic drift cannot explain the speed of fixation of albinism in this case, suggesting that other evolutionary forces are involved, possibly based on one or more of the multiple advantages of the loss of pigmentation we have discovered in this study.

## Acknowledgements

This paper is dedicated to the memory of Daniel Wu Fong (1954-2025). Daphne Soares and Matthew Niemiller are thanked for providing *Typhlichthys subterraneus* and *Amblyopsis spelaea* samples. Branko Jalžić, Jana Bedek, Alen Kirin, Ana Komerički, Marko Lukić, Kazimir Miculinić, Martina Pavlek, and other members of Croatian Biospeleological Society are thanked for field assistance and for providing samples.

