## Supplemental table 1 for "A common mechanism and multiple advantages of pigment loss underlie the convergent evolution of albinism in cave animals"

One way Analysis of Variance was performed separately for two HPLC runs used to quantify the dopamine (DA), DOPAC and the ratio of DOPAC/DA in brains of *Astyanax mexicanus* surface fish (SF), four cavefish populations from the Pachón (PA), Tinaja (TI), Los Sabinos (LS) and Molino (MO) caves, two hybrid groups, the F1 generation of Pachón cavefish surface fish cross, and the 4^th^ generation of the hybrid offspring selected for albino phenotype but fully developed eyes (AE). All Pairwise Multiple Comparison Procedures followed the Holm-Sidak method. Unadjusted p values are given in the table for all pairwise comparisons and the overall significance level was set at < 0.001.

DA, measured on September 2016.

| GROUP | SF | PA | TI | LS | MO | F1 |
| --- | --- | --- | --- | --- | --- | --- |
| SF |  | 0.00158 | 0.00299 | 0.00157 | 0.730 | 0.858 |
| PA |  |  | 0.958 | 0.801 | 0.00178 | 9.96 x 10^-4^ |
| TI |  |  |  | 0.856 | 0.00301 | 0.00196 |
| LS |  |  |  |  | 0.00192 | 9.56 x 10^-4^ |
| MO |  |  |  |  |  | 0.850 |
| F1 |  |  |  |  |  |  |

DOPAC, measured on September 2016.

| GROUP | SF | PA | TI | LS | MO | F1 |
| --- | --- | --- | --- | --- | --- | --- |
| SF |  | 8.65 x 10^-5^ | 0.0346 | 2.52 x 10^-7^ | 0.305 | 0.974 |
| PA |  |  | 0.0659 | 0.148 | 0.00514 | 7.90 x 10^-5^ |
| TI |  |  |  | 0.00163 | 0.304 | 0.0324 |
| LS |  |  |  |  | 5.95 x 10^-5^ | 2.29 x 10^-7^ |
| MO |  |  |  |  |  | 0.291 |
| F1 |  |  |  |  |  |  |

DOPAC/DA measured on September 2016.

| GROUP | SF | PA | TI | LS | MO | F1 |
| --- | --- | --- | --- | --- | --- | --- |
| SF |  | 0.0169 | 0.845 | 1.41 x 10^-5^ | 0.00364 | 0.964 |
| PA |  |  | 0.0476 | 0.0350 | 0.393 | 0.0187 |
| TI |  |  |  | 1.56 x 10^-4^ | 0.0111 | 0.876 |
| LS |  |  |  |  | 0.312 | 1.61 x 10^-5^ |
| MO |  |  |  |  |  | 0.00402 |
| F1 |  |  |  |  |  |  |

DA, measured on April 2016.

| GROUP | SF | PA | TI | AE |
| --- | --- | --- | --- | --- |
| SF |  | 4.89 x 10^-11^ | 0.000000245 | 0.000795 |
| PA |  |  | 0.00860 | 0.000371 |
| TI |  |  |  | 0.106 |
| AE |  |  |  |  |

DOPAC, measured on April 2016.

| GROUP | SF | PA | TI | AE |
| --- | --- | --- | --- | --- |
| SF |  | 7.969E-012 | 0.0000517 | 0.0000784 |
| PA |  |  | 0.00000413 | 0.000355 |
| TI |  |  |  | 0.566 |
| AE |  |  |  |  |

DOPAC/DA, measured on April 2016.

| GROUP | SF | PA | TI | AE |
| --- | --- | --- | --- | --- |
| SF |  | 0.0000212 | 0.297 | 0.00291 |
| PA |  |  | 0.000571 | 0.498 |
| TI |  |  |  | 0.0264 |
| AE |  |  |  |  |
